# Allosteric substrate activation of SAMHD1 shapes deoxynucleotide triphosphate imbalances by interconnecting the depletion and biosynthesis of different dNTPs

**DOI:** 10.1101/2023.11.14.567083

**Authors:** Claudia McCown, Corey H. Yu, Dmitri N. Ivanov

## Abstract

SAMHD1 is a dNTPase that impedes replication of HIV-1 in myeloid cells and resting T lymphocytes. Here we elucidate the substrate activation mechanism of SAMHD1 that depends on dNTP binding at allosteric sites and the concomitant tetramerization of the enzyme. The study reveals that SAMHD1 activation involves an inactive tetrameric intermediate with partial occupancy of the allosteric sites. The equilibrium between the inactive and active tetrameric states, which is coupled to cooperative binding/dissociation of at least two allosteric dNTP ligands, controls the dNTPase activity of the enzyme, which, in addition, depends on the identity of the dNTPs occupying the four allosteric sites of the active tetramer. We show how such allosteric regulation determines deoxynucleotide triphosphate levels established in the dynamic equilibria between dNTP production and SAMHD1-catalyzed depletion. Notably, the mechanism enables a distinctive functionality of SAMHD1, which we call facilitated dNTP depletion, whereby elevated biosynthesis of some dNTPs results in more efficient depletion of others. The regulatory relationship between the biosynthesis and depletion of different dNTPs sheds light on the emerging role of SAMHD1 in the biology of dNTP homeostasis with implications for HIV/AIDS, innate antiviral immunity, T cell disorders, telomere maintenance and therapeutic efficacy of nucleoside analogs.

## INTRODUCTION

HD-domain dNTPases catalyze hydrolysis of deoxynucleotide triphosphates to dephosphorylated deoxynucleosides and inorganic triphosphate^1-5^. This enzymatic reaction is thought to enable controlled depletion of cellular dNTP reservoirs, because the reentry of dephosphorylated deoxynucleosides to the dNTP pools is limited by the deoxynucleoside salvage pathway (Fig 1A). The human HD-domain dNTPase, SAMHD1, impedes the ability of HIV-1 and related retroviruses to infect myeloid cells and resting T-lymphocytes and is specifically targeted for degradation by the Vpx protein encoded by the HIV-2 and SIV^6-8^. The antiretroviral function of SAMHD1 brought to light a mechanism of intrinsic cellular antiviral immunity that depends on the enzymatic degradation of deoxynucleotide triphosphates. Subsequent studies revealed that SAMHD1 can also limit infectivity and pathogenesis of DNA viruses^9-14^, and that deoxynucleotide depletion is similarly used for antiviral defense by bacteria^15,16^. Notably, SAMHD1-like dNTPases are ubiquitous in prokaryotic genomes throughout the evolutionary tree^17^.

**Figure 1.**
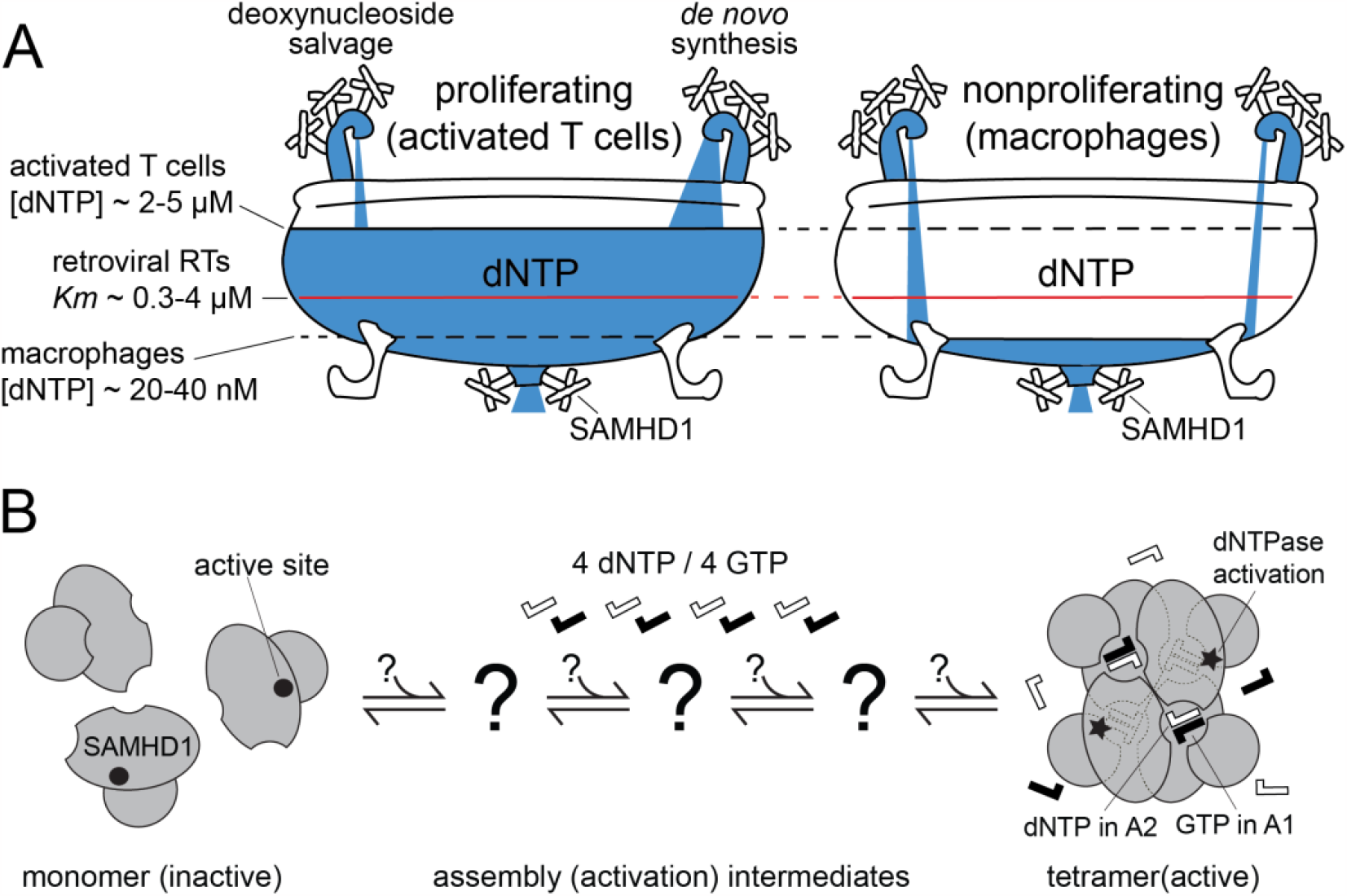
SAMHD1 is a key regulator of dNTP homeostasis. (A) Cellular dNTP levels are established in a dynamic equilibrium between dNTP biosynthesis by the de-novo and salvage pathways and dNTP depletion by SAMHD1 and DNA polymerases. SAMHD1 limits dNTP concentrations and impedes retroviral replication in nonproliferating immune cells. (B) Allosteric activation of SAMHD1 requires binding of GTP and dNTP ligands at the allosteric sites A1 and A2, which promotes assembly of the enzyme into a catalytically active tetramer^44-50^. Here we investigate the mechanism of this unusual, tetramerization-dependent activation.

It is well established that the rate of the de novo dNTP biosynthesis varies greatly at different stages of the cell cycle and is downregulated in nonproliferating cells^18-21^. For example, dNTP concentrations in macrophages are in the mid-nanomolar range, which is approximately 100-fold lower than the concentrations measured in activated T lymphocytes^22,23^. SAMHD1 activity is a critical limiting factor of dNTP availability in nonproliferating immune cells, and SAMHD1-mediated decrease in the dNTP levels in myeloid cells correlates with the inability of lentiviruses to undergo reverse transcription ^5,8,24-26^. However, even in nonproliferating cells active dNTP synthesis is required for DNA repair and mitochondrial biogenesis^27-31^, and it is not clear how myeloid cells can inhibit reverse transcription of the 10 kb retroviral genome but maintain replication of the 15 kb mitochondrial DNA. Notably, transient modulation of SAMHD1 activity in macrophages, triggered by various cellular signaling events, makes them susceptible to HIV infection^32-38^. These observations suggest that mechanisms of temporal or compartmental control of dNTP availability may impact HIV-1 infection of monocytes and macrophages in vivo, where HIV-1 infected myeloid cells contribute to HIV pathogenesis and latency^39-42^. Little is currently known about temporal or compartmental regulation of dNTP reservoirs in nonproliferating cells because experimental tools for monitoring distinct states of dNTP homeostasis remain limited despite decades of cellular dNTP measurements^43^.

To gain further insight into antiviral modalities of dNTP metabolism we turned to the allosteric regulation of dNTP hydrolysis by SAMHD1. SAMHD1 is activated by the transient assembly of the enzyme into the catalytically active tetramer, which is promoted in turn by the binding of allosteric nucleotide activators at two adjacent allosteric sites, A1 and A2^44-50^ (Fig. 1B). The A1 site is selective for guanine nucleotides, GTP or dGTP, whereas the A2 site is dNTP specific but can accommodate any nucleobase. Many questions remain about the exact contribution of this mechanism to the regulation of dNTP homeostasis and HIV-1 restriction by SAMHD1. Substrate activation is a common mechanism of establishing stable cellular concentrations of metabolites, but activation of metabolic enzymes by transient oligomerization is unusual. The rate of protein assembly into functional complexes is inherently limited by the slow diffusion of large polypeptides^51^. It takes a SAMHD1 monomer a relatively long time to find three other mates, and, as a necessary consequence, the dissociation of SAMHD1 tetramers at physiological conditions is also very slow. Indeed, it has been shown that SAMHD1 tetramers can persist for more than an hour after dNTPs have been depleted by the enzyme^52-54^. The 1000-fold mismatch between the *k*_*cat*_ (∼1-5 sec^-1^) and the dissociation rate (∼10^-3^ sec^-1^) of the catalytically active SAMHD1 tetramer is problematic from the point of view of control theory, because regulatory devices, whose response time is slower than the intrinsic variability of the processes that they regulate, are prone to oscillations, overshooting and overall instability. The role of this mechanism in HIV restriction is also puzzling. SAMHD1 tetramerization requires micromolar concentrations of dNTPs, so it is not clear whether and how this mechanism contributes to the regulation of the mid-nanomolar dNTP concentrations observed in nonproliferating immune cells, where the HIV-1 restriction activity of SAMHD1 is manifested^54,55^.

Here, we show that SAMHD1 activation is a two-step process, consisting of tetramer assembly and tetramer activation, which explains how the enzyme can rapidly react to changing dNTP levels despite its inherently slow assembly and disassembly. Most notably, the mechanism gives rise to a distinctive functionality that we call facilitated dNTP depletion. Facilitated dNTP depletion enables more efficient depletion of some nucleotides when biosynthesis of certain others is elevated. The findings offer insight into the antiviral effects of dNTP imbalances and several other recent observations implicating SAMHD1 in the biology of dNTP homeostasis.

## RESULTS

### The overall catalytic efficiency, but not substrate specificity, of SAMHD1 depends on the identity of the dNTPs occupying the A2 allosteric sites

SAMHD1 accommodates all four deoxynucleotide triphosphates as substrates and allosteric activators. In this study, we aimed to determine how the catalytic activity of SAMHD1 is affected by the identity of the allosteric dNTP ligands. To monitor the dNTPase activity of SAMHD1, we leveraged the well-resolved NMR signals of dNTP nucleobases that are sensitive to the hydrolysis of the triphosphate group (Fig. 2A)^56^. The NMR assay offers simplicity, a real-time readout, and the ability to independently monitor the hydrolysis of individual deoxynucleotides in a dNTP mixture. The method is remarkably robust in determining the *k*_*cat*_ and *K*_*m*_ values because the *K*_*m*_ values of SAMHD1 for all dNTP substrates far exceed the NMR detection sensitivity (Methods). *k*_*cat*_ /*K*_*m*_ values were used to quantify the catalytic efficiency of the enzyme.

**Figure 2.**
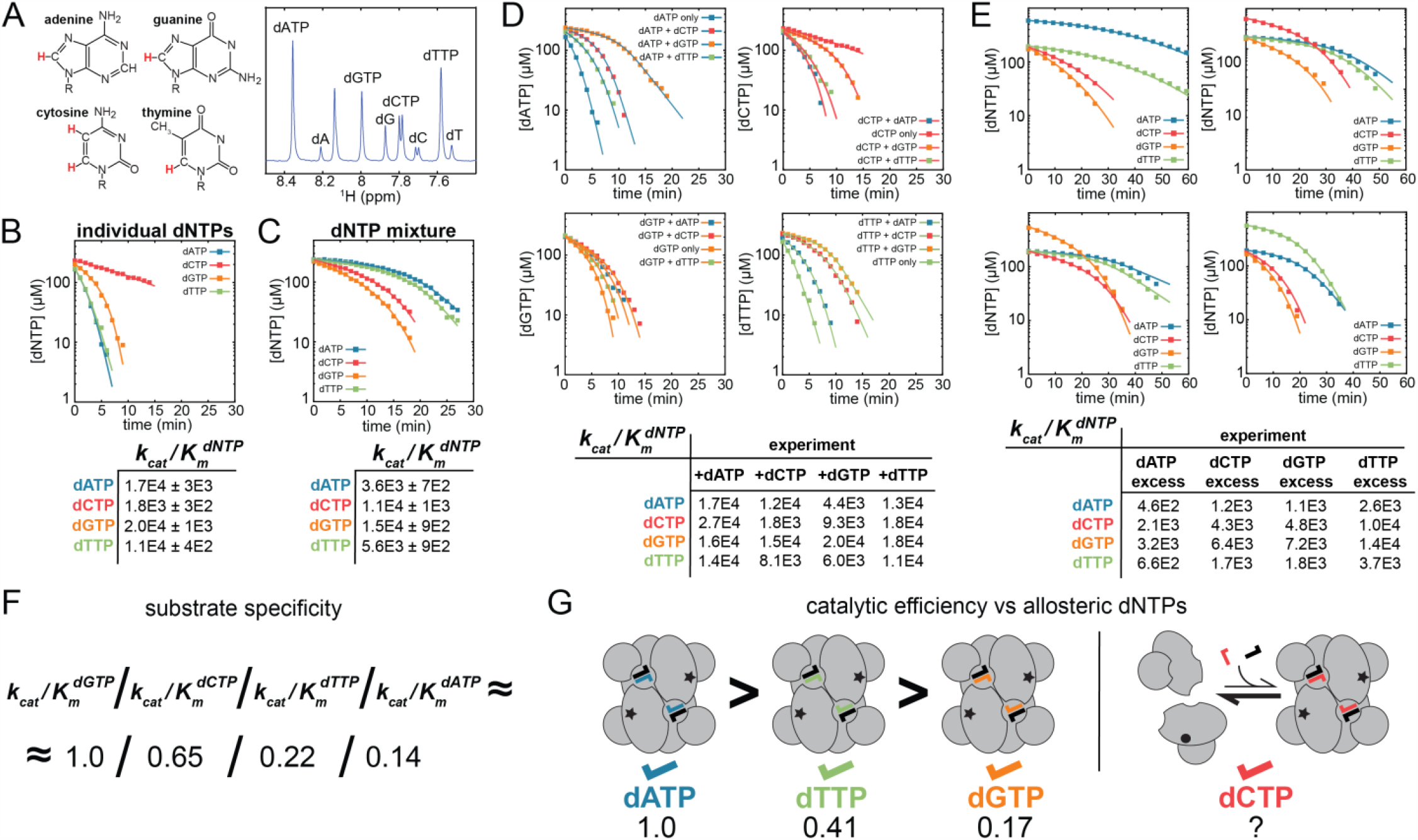
NMR-monitored dNTPase assays reveal how allosteric ligand identity affects catalytic activity. (A) Non-overlapping ^1^H NMR signals of the aromatic nucleobase protons (highlighted in red) enable real-time monitoring of dNTP hydrolysis in samples containing several substrates. (B-E) k_cat_/K_m_ values for each dNTP substrate were determined by fitting reaction progress curves to the Michaelis-Menten model (see Methods). All reactions contained 500 μM of GTP and 1 μM of SAMHD1_114-626_. (B,C) Hydrolysis efficiencies of the four substrates (k_cat_/K_m_) are markedly different in reactions performed individually (B) or in the equimolar dNTP mixture (C). (D) Reactions performed in all possible combinations of two substrates reveal how the competition of two different dNTPs for the allosteric sites impacts the catalytic efficiency of the active site for each substrate. (E) Similarly, reactions initiated in imbalanced dNTP mixtures demonstrate the dependence of the catalytic efficiency on the identity of the allosteric dNTP ligands. (F) The ratio of k_cat_/K_m_ values for different dNTP substrates remains approximately constant in all reactions containing more than one substrate (C,D,E). This finding reveals that the specificity of the active sited does not depend on the identity of the dNTP ligands occupying the allosteric sites. (G) In contrast, the overall catalytic efficiency varies significantly depending on the occupancy of the allosteric sites by different dNTP ligands. For example, when the same dNTP ligand occupies all four allosteric sites (B), the dATP>dTTP>dGTP relationship is observed. The ability of dCTP to activate SAMHD1 on its own is significantly lower than that of other dNTPs.

First, we determined *k*_*cat*_ /*K*_*m*_ values for dNTPase reactions containing individual deoxynucleotides (Fig. 2B). In this set of experiments, the same dNTP occupies all four A2 sites within the SAMHD1 tetramer, thereby activating its own hydrolysis. Reactions initiated with two distinct dNTP concentrations, 250 μM and 500 μM, yielded similar *k*_*cat*_ /*K*_*m*_ values with good overall agreement between the experimental progress curves and the predictions of the Michaelis-Menten model. (Supplemental Fig. S1ABCD). The relatively low catalytic efficiency of dCTP hydrolysis by SAMHD1 compared to other dNTPs was a notable finding in this set of experiments.

We then measured *k*_*cat*_ /*K*_*m*_ for each deoxynucleotide in an equimolar mixture containing 250 μM of each dNTP (Fig. 2C). We observed that the *k*_*cat*_ /*K*_*m*_ of dGTP hydrolysis was only modestly reduced in the mixture, but the hydrolysis efficiencies of the other three dNTPs exhibited marked differences. In contrast to the single-dNTP reactions, the dCTP *k*_*cat*_ /*K*_*m*_ increased more than 6-fold in the mixture, whereas the dATP and dTTP *k*_*cat*_ /*K*_*m*_ were reduced 4.7-fold and 2-fold, respectively. The observed changes in the *k*_*cat*_ /*K*_*m*_ values do not stem from the competition among different dNTP substrates for the active site; this factor was explicitly accounted for in the fitting procedure (Methods). Instead, the *k*_*cat*_ /*K*_*m*_ variations signify that different dNTP occupancies of the A2 sites alter the catalytic efficiency of the enzyme.

To further investigate this, we measured *k*_*cat*_ /*K*_*m*_ in all possible pairwise combinations containing 250 μM of each dNTP (Fig. 2D). These results offer additional insight into patterns of SAMHD1 activation by different allosteric dNTPs. For example, dGTP hydrolysis is not significantly altered by the addition of dATP, dTTP or dCTP, which suggests that dGTP is not efficiently displaced from the A2 sites by the other dNTPs in equimolar mixtures. The finding indicates that dGTP has the highest affinity for the A2 sites among the four dNTPs. In contrast, depletion of dATP and dTTP is significantly less efficient in the dGTP-containing mixtures than for pure dATP and dTTP, which indicates that the catalytic efficiency of SAMHD1 containing dGTP in the A2 sites is lower than that for the dATP-or dTTP-activated tetramers. In addition, the *k*_*cat*_ /*K*_*m*_ of dCTP hydrolysis is significantly enhanced in all mixtures, which indicates that dCTP is a poor allosteric activator of SAMHD1 on its own but is a perfectly good substrate for the activated enzyme.

Finally, we performed *k*_*cat*_ /*K*_*m*_ measurements in four imbalanced dNTP mixtures containing 600 μM of one dNTP and 200 μM each of the other three (Fig. 2E). Despite considerable variations in overall catalytic efficiency across different experiments, the ratios of *k*_*cat*_ /*K*_*m*_ values for the four dNTPs demonstrated a consistent pattern in all samples: dGTP/dCTP/dTTP/dATP = 1.0/0.65/0.22/0.14 (Fig 2F). The same approximate ratios of the *k*_*cat*_ /*K*_*m*_ values were also observed in the pairwise mixtures. These observations strongly suggest that the specificity of the enzyme’s active site for dNTP substrates is not affected by the identity of the allosteric dNTP activators occupying the A2 sites. Therefore, one can use these *k*_*cat*_ /*K*_*m*_ ratios to directly compare overall catalytic efficiencies of tetramers in experiments with individual dNTPs, where the same dNTP ligand is bound at all four A2 sites. This analysis yields 1.0/0.41/0.17 activity ratios for the dATP-, dTTP- and dGTP-activated tetramers, assuming that the enzyme is fully activated at the high dNTP concentrations used in these experiments. The poor ability of dCTP to promote SAMHD1 activation is unique among the four dNTPs because even at the highest tested dCTP concentrations SAMHD1 was not fully active (Fig. 2G).

### *EC*_*50*_ values for dNTP-dependent tetramerization of SAMHD1 are in the low micromolar range

To better understand the tetramerization-dependent activation of SAMHD1 we developed a FRET assay that allows monitoring of SAMHD1 tetramerization independently of the dNTPase activity. To this end, we labeled the N-terminus of SAMHD1 with either donor (AF488) or acceptor (AF594) fluorophores using sortase-catalyzed transpeptidation (Fig. 3A and Methods)^57^. In the FRET assay, SAMHD1 tetramerization manifests as a dNTP-dependent increase in the FRET signal in a sample containing an equimolar mixture of the AF488- and AF594-labeled proteins (Fig. 3B). For example, adding dNTPs to the WT SAMHD1 sample results in an initial increase in FRET signal intensity, which subsequently decays as the enzyme hydrolyzes the dNTPs (Fig. 3CD). Notably, in this series of experiments, dCTP stands out due to the markedly slower decay of its FRET signal compared to the other dNTPs (Fig. 3CD).

**Figure 3.**
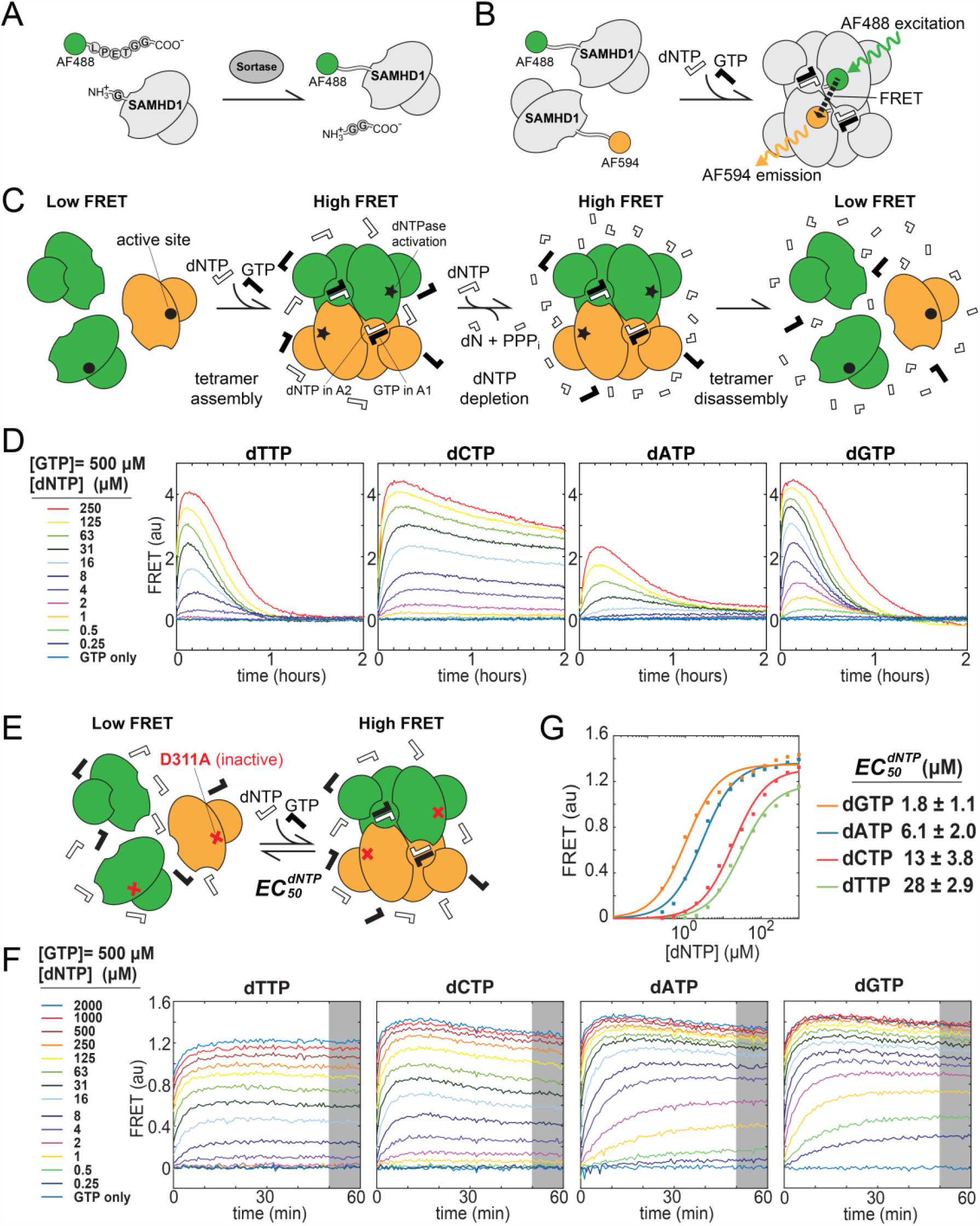
FRET studies of SAMHD1 tetramerization and its dependence on dNTP concentrations. (A) N-terminal labeling of SAMHD1 with fluorophores using sortase. (B) Proximity of N-termini of two neighboring monomers within the tetramer enables monitoring of SAMHD1 tetramerization by FRET. (C) The tetramerization of WT SAMHD1, initiated by the addition of GTP and dNTP, is a reversible process, with tetramers dissociating upon hydrolysis of dNTP. (D) This transient behavior is captured by the rise and fall of the FRET signal over time. (E) In contrast, a stable equilibrium is established for the D311A SAMHD1 mutant, which is not catalytically active. (F,G) Dependence of the equilibrium FRET signal on dNTP concentration can be used to determine the *EC*_*50*_ values for dNTP-dependent tetramerization of SAMHD1.

To determine the *EC*_*50*_ values for the dNTP-dependent tetramerization of SAMHD1 we performed dNTP titrations with a catalytically inactive D311A mutant of SAMHD1. The D311A mutation disrupts coordination of the catalytic metal ion in the active site but does not affect the tetramerization of the protein^56^. The catalytic activity of the WT subunits is not affected by the presence of the D311A subunits within SAMHD1 tetramers formed in WT/D311A mixtures, which confirms that the D311A mutation does not affect the allosteric functionality of the protein (Supplemental Fig. S2). Upon adding dNTPs, the FRET signal of the D311A mutant intensifies and stabilizes at a signal intensity dependent on the dNTP concentration (Fig 3F). This dependence can be used to measure the tetramerization *EC*_*50*_ values for different dNTPs (Fig. 3G). The *EC*_*50*_ values determined by this method fall in the low micromolar range and agree with the previously reported *EC*_*50*_ values for allosteric activation of SAMHD1^54,55^. dGTP has the lowest apparent *EC*_*50*_, which is consistent with its ability to outcompete other dNTPs for the A2 sites as revealed by the NMR dNTPase experiments (Fig. 2D).

### SAMHD1 activation involves a tetrameric intermediate with partially loaded A2 sites

As we explain briefly in the introduction, an activation mechanism that depends on transient assembly and disassembly of protein oligomers presents a kinetic problem and limits the ability of an enzyme to rapidly react to changing dNTP concentrations. On the other hand, the remarkably slow dissociation of the catalytically active SAMHD1 tetramer was proposed to be critical for the ability of SAMHD1 to deplete cellular dNTP pools and to restrict retroviral replication^53^.

To evaluate the contribution of tetramer half-life to SAMHD1 function we used the FRET assay to accurately measure tetramer dissociation rates and their dependence on the identity of the dNTP ligand occupying the A2 allosteric site. To this end, the FRET-active D311A SAMHD1 sample (an equimolar mixture containing 100 nM each of the AF488- and AF594-labeled D311A SAMHD1) was first incubated with 200 μM of dNTP and 50 μM of GTP until the FRET signal reached its maximum equilibrium value. After that 25x excess (2.5 μM) of unlabeled SAMHD1 protein was added to the samples and the subsequent decay of the FRET signal intensity was followed over time (Fig. 4A). The experiment yielded dramatically different results depending on whether the wild-type SAMHD1 (Fig. 4B) or the inactive D311A mutant (Fig. 4C) was used as the excess unlabeled protein. In the experiments with the catalytically active WT SAMHD1, which rapidly degrades dNTPs present in the sample, the FRET signals decayed as expected and the apparent decay rates were different for different dNTPs (Fig. 4B). In contrast, when the dNTPase-inactive D311A mutant was used, no significant decay in the FRET intensity was observed after several hours (Fig. 4C).

**Figure 4.**
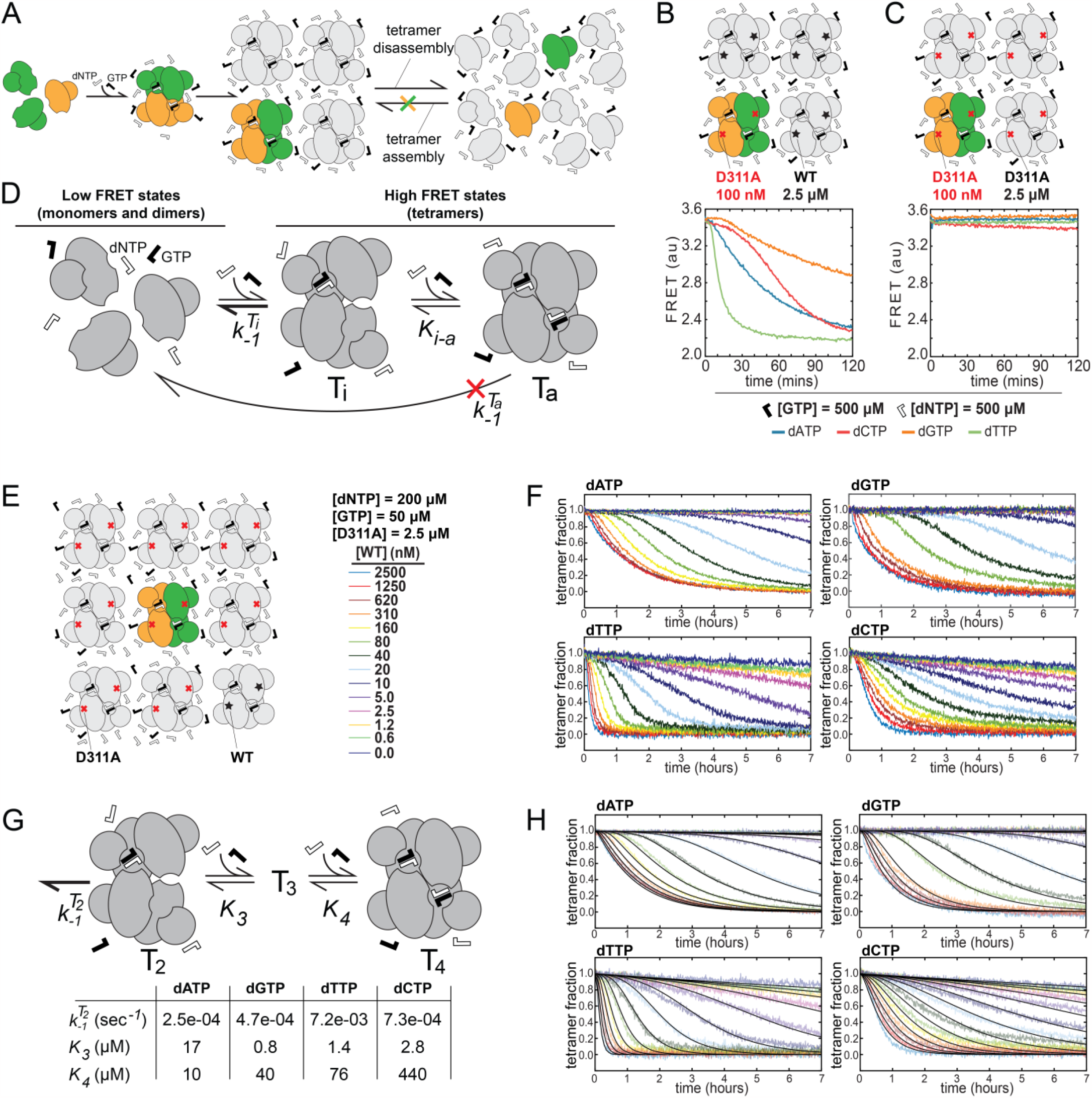
Discovery and characterization of a SAMHD1 activation intermediate with incomplete occupancy of the allosteric sites. (A) Tetramer dissociation rates can be measured by mixing preassembled, high-FRET D311A SAMHD1 tetramers with a large excess of unlabeled protein. (B,C) FRET decay rates are markedly different when high-FRET tetramers are mixed with unlabeled WT SAMHD1 (B) or with unlabeled D311A SAMHD1 (C). (D) FRET decay data establish that a tetrameric state with incomplete occupancy of allosteric sites is a required intermediate in the activation pathway. (E,F) Dilution of high-FRET tetramers into a mixture of the unlabeled D311A and WT SAMHD1 (E) offers additional insight into the mechanism of SAMHD1 assembly/dissociation and reveals notable differences between different dNTPs (F). (G,H) The data suggest that the equilibrium between the partially-loaded and fully-loaded SAMHD1 tetramers involves binding of two dNTP ligands (G) and the model yields good overall agreement between the theory (solid black lines) and experiment (faded color lines) (H).

The strong dependence of the apparent FRET decay rate on the free dNTP concentration reveals that formation of a high-FRET SAMHD1 tetramer takes place prior to full loading of the SAMHD1 tetramer with the allosteric dNTP ligands (Fig. 4D). The findings establish that the tetramerization-dependent activation of SAMHD1 involves a tetrameric intermediate (T_i_ in Fig. 4D), in which only some of the A2 sites are occupied by dNTPs; conversely, the dissociation of the fully dNTP loaded, enzymatically-active SAMHD1 tetramer (T_a_ in Fig. 4D) must proceed through this partially dNTP loaded intermediate T_i_. As a result, when dNTP concentrations are high, the equilibrium between T_i_ and T_a_ is shifted to the right and very little tetramer dissociation is observed. When dNTP concentrations are low and the equilibrium is shifted to the left, the FRET decay rate equals *k*_*-1*_^*Ti*^ (T_i_ dissociation rate) times T_i_ concentration.

### Affinity and cooperativity of allosteric dNTP binding control the equilibrium between the partially dNTP loaded and fully dNTP loaded tetrameric states

To better understand how the equilibrium between distinct tetrameric states depends on dNTP concentrations we slightly modified our experimental strategy for monitoring tetramer dissociation. In this series of experiments fluorescently labeled D311A SAMHD1 was first assembled into FRET-active tetramers and then diluted into large excess of unlabeled protein. This time, the unlabeled protein contained 2.5 μM of the D311A mutant and increasing amounts of the wild-type enzyme (Fig. 4E). The resulting data-rich datasets display notable differences in the curvature and distribution of the individual FRET decay curves in the data series acquired for different deoxynucleotides (Fig. 4F).

FRET decay curves can be numerically simulated in MATLAB using the theoretical model in Fig. 4D (Supplemental Fig. S3C and Supplemental Methods). This analysis is built upon three key assumptions: first, the tetramer dissociation necessarily proceeds through the partially loaded tetrameric intermediate T_i_, which is corroborated by the data in Fig. 4BC; second, only the fully dNTP-loaded tetramer has catalytic activity — a point to be elaborated upon in the following section; finally, the equilibration among different tetrameric states occurs substantially faster than the tetramer dissociation process.

Numerical analysis delineates how the FRET decay datasets are influenced by the three key model parameters: the tetramer dissociation rate *k*_*-1*_, the dissociation constant *K*_*i-a*_,which describes the dNTP-dependent equilibrium between the T_i_ and T_a_ tetrameric states, and the Hill coefficient *n* pertinent to this equilibrium (Supplemental Fig. S3ABCD).

Analysis of the dATP data series revealed that excellent correspondence between theory and experiment can be achieved using Hill coefficient values in the range of 1.6-1.7, implying that binding of at least two dNTP ligands accompanies the transition between the T_i_ and T_a_ states. Structural considerations suggest that binding of two allosteric dNTPs is required for the assembly of two SAMHD1 dimers into a tetramer, whereas binding of two more promotes the transition from T_i_ to T_a_ (see Discussion). The equilibrium between different tetrameric states can therefore be described by the Adair equation with two binding constants *K*_*3*_ and *K*_*4*_ representing binding of the third and fourth allosteric dNTP, respectively (Fig. 4G and Supplemental Methods).

The nonlinear least-squares fitting of the *k*_*-1*_, *K*_*3*_ and *K*_*4*_ parameter values of the model yields good overall agreement between theory and experiment for all four dNTPs, although some systematic deviations are noticeable in the pyrimidine datasets (Fig. 4H). The tetramer dissociation rate *k*_*-1*_ can be determined with higher degree of confidence because of the dominant effect of the *k*_*-1*_ on the FRET decays at high WT SAMHD1 concentrations. *K*_*3*_ and *K*_*4*_ binding constants are less well constrained by the model individually because their absolute values depend on the accurate quantification of the active enzyme and the precision of the *k*_*cat*_ /*K*_*m*_ measurements. However, their ratio *K*_*3*_ */K*_*4*_, which determines the curvature and the distribution of the individual FRET decay curves within one experimental series, reveals positive cooperativity of dATP binding to the allosteric sites. In contrast, dCTP binding is strongly anti-cooperative, which, as we show in the next section, explains the poor ability of dCTP to promote SAMHD1 activation.

### The dNTP-dependent equilibrium between distinct tetrameric states regulates the dNTPase activity of SAMHD1

To further explore how the equilibrium between distinct tetrameric states impacts the dNTPase activity of SAMHD1 we sought to develop a method for monitoring dNTP hydrolysis at low dNTP concentrations where the allosteric inactivation of the enzyme becomes apparent. Limited sensitivity of the NMR dNTPase assay makes it unsuitable for such experiments. Instead, we turned to dNTP quantification by LC-MS, the sensitivity of which is greatly superior to that of NMR. Briefly, SAMHD1 was incubated with dNTPs for increasing time intervals, reactions were then stopped by adding acetic acid to a 10% final concentration and the residual dNTPs in the sample were quantified by LC-MS as described in the Methods section. The LC-MS detection enables accurate monitoring of the dNTPase progress curves all the way to completion, where the reaction effectively stops because of allosteric inactivation of SAMHD1 (Fig. 5AB).

**Figure 5.**
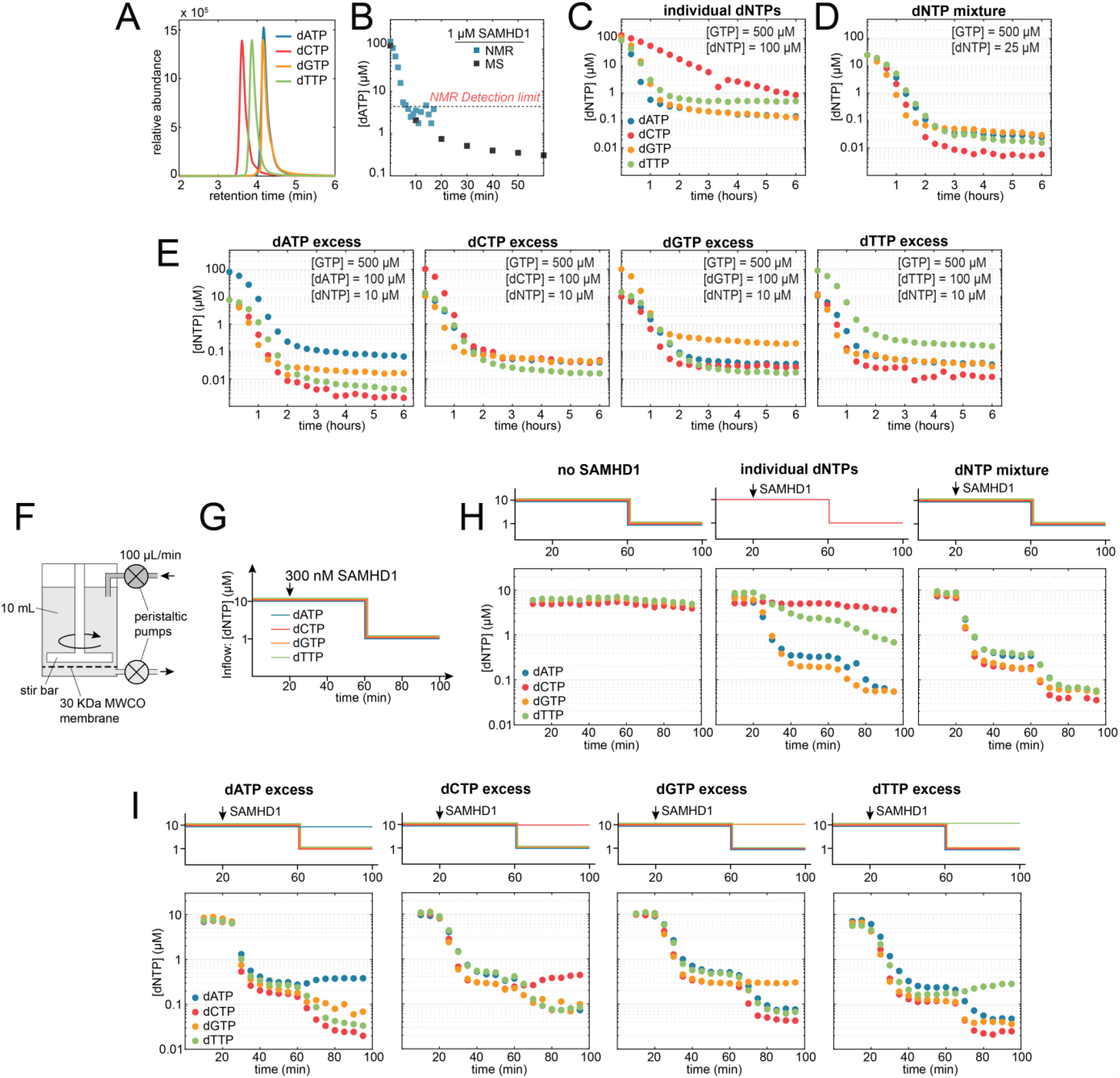
LC-MS analysis of dNTP depletion reveals a regulatory relationship between the depletion and biosynthesis of different dNTPs. (A,B) High sensitivity of dNTP quantification by LC-MS (A) enables accurate measurements of residual dNTP concentrations in the hydrolysis reactions catalyzed by SAMHD1 (B). (C,D) dNTP depletion reactions performed with individual dNTPs reveal poor ability of dCTP to promote its own hydrolysis (C). dNTPs are depleted to significantly lower final concentrations when the hydrolysis reaction is initiated in an equimolar mixture of four dNTPs (D). (E) Reactions initiated in imbalanced dNTP mixtures reveal that dATP excess promotes pyrimidine depletion to <10 nM final concentrations. (F) Dynamic dNTP equilibria established by SAMHD1 were investigated in a continuous flow reaction chamber. (G) Experimental schematic: delivery of dNTP substrates into the chamber was controlled by varying dNTP concentrations in the inflow. (H) Reactions performed with individual dNTPs (center panel) and in an equimolar mixture (right panel) reveal critical dependence of pyrimidine dNTP depletion on purine dNTP availability. (I) Effects of dNTP supply imbalances. Elevated supply of dATP further decreases pyrimidine dNTP concentrations established in the reactions.

Consistent with the findings from the NMR-monitored experiments, hydrolysis reactions performed with individual deoxynucleotide triphosphates reveal the markedly low efficiency of dCTP depletion by SAMHD1 (Fig. 5C). In contrast, in the mixtures containing other dNTPs, dCTP hydrolysis is more robust than that of dATP and dTTP (Fig. 2 and Fig. 5D), which establishes that dCTP is a good substrate but a poor allosteric activator of SAMHD1. The weak activation of SAMHD1 by dCTP is notable, given that the efficacy of dCTP in promoting the formation of the high-FRET tetramer is on par with other nucleotides (Fig. 3G). These observations indicate that the tetrameric intermediate with partial dNTP occupancy of the A2 sites (T_i_ in Fig. 4D) is not catalytically active and that full loading of the allosteric sites with dNTP ligands (T_a_ in Fig. 4DG) is required for the dNTPase activity. This requirement explains how strong negative cooperativity of dCTP binding to the A2 sites (Fig. 4GH) impairs the ability of dCTP to activate catalysis, without affecting its ability to promote assembly of the high-FRET SAMHD1 tetramer.

Dependence of dNTPase activity on full occupancy of A2 sites can also be established for other deoxynucleotides. Owing to the slow tetramer dissociation rate *k*_*-1*_, the SAMHD1 activation model in Fig. 4D predicts depletion to significantly lower dNTP concentrations if activity of the T_i_ tetramer is comparable to that of T_a_. Comparison of the model predictions to experimental dATP depletion outcomes demonstrates that full occupancy of allosteric sites is indeed required for dNTPase activation (Supplemental Fig. S4B).

The data establish that SAMHD1 tetramerization is required but not sufficient for the activation of catalysis. The outcomes of dNTP depletion reactions reveal that the dNTPase activity is controlled by the binding and dissociation of allosteric dNTP ligands to the preformed SAMHD1 tetramer, rather than by transient changes in the SAMHD1 oligomerization states. This mechanism circumvents the kinetic problem posed by the inherently slow assembly and disassembly of protein oligomers because binding of small molecules and conformational changes within protein complexes are not limited by the low diffusion rates of large macromolecules.

### Facilitated dNTP depletion: efficient pyrimidine depletion depends on purine availability

Comparing the depletion outcomes for individual nucleotides with those for the equimolar dNTP mixture reveals another important functional consequence of the substrate activation mechanism described here. SAMHD1 can accommodate any dNTP in both its active site and its A2 allosteric sites, which raises the question of whether and how different dNTPs influence each other’s depletion. In experiments described above, we establish that the identity of allosteric dNTP ligands affects the overall catalytic efficiency of SAMHD1 and that there are notable differences in the affinities and cooperativities of allosteric site loading by different dNTPs. Depletion reactions initiated at 100 μM of individual dNTPs are notably less efficient for pyrimidine dNTPs than for purine dNTPs, which is expected given the less potent affinity/cooperativity of pyrimidine loading of the A2 sites. Pyrimidine depletion becomes remarkably more robust in an equimolar mixture containing 25 μM of each dNTP. These observations establish that different dNTPs can indeed facilitate each other’s hydrolysis, because efficient depletion of pyrimidine dNTPs strongly depends on purine dNTP availability. The effect is less pronounced for dGTP and dATP hydrolysis, but they are also depleted to lower final concentrations in the mixed dNTP sample, which suggests that mixed loading of the four A2 sites with different dNTP ligands may play an important role in facilitated depletion of all dNTPs by SAMHD1.

Further insight into the interplay between the hydrolysis and availability of different dNTPs emerges from the reactions initiated in imbalanced dNTP mixtures, in which one dNTP is present at a 10-fold excess over the others (Fig. 5E). In these reactions, the initial excess of one nucleotide is maintained throughout the experiment, except for the dCTP excess, which is eliminated during the depletion reaction. Excess of dCTP or dTTP has little effect on the outcomes for the other three dNTPs, which may be explained by their limited ability to outcompete purine dNTPs for the allosteric sites even at elevated concentrations. Most notable is the effect of dATP excess, which results in additional significant reduction in final pyrimidine concentrations. Pyrimidines are depleted to concentrations two to three orders of magnitude lower in the reactions with dATP excess than in the reactions containing dCTP or dTTP alone, which demonstrates that dATP availability is a major determinant of pyrimidine depletion in dNTP mixtures.

### Deoxynucleotide triphosphate homeostasis is shaped by the interdependence between the depletion and biosynthesis of different dNTPs

In cells, dNTP levels are established in a dynamic equilibrium between dNTP production by the de-novo and salvage synthesis and dNTP depletion by SAMHD1 and DNA polymerases (Fig. 1A). To investigate the contribution of SAMHD1 to these dynamic equilibria we constructed a continuous-flow reaction chamber by modifying a pressurized protein concentrator (Fig. 5F). The concentrator was equipped with an input line, through which dNTP solution of a desired composition was delivered into the reaction chamber, whereas dNTP concentrations within the chamber were measured at different time points by performing LC-MS analysis of the outflow. The 30 kDa molecular weight cutoff membrane retained SAMHD1 within the chamber but did not impede the outflow of dNTPs. Experiments were performed by filling the reaction chamber with 10 mL solution of either individual dNTPs at 10 μM or an equimolar mixture at 10 μM each. Continuous flow of the same dNTP solution was then established through the chamber at 100 μL/min (Fig. 5GH). After 15 minutes SAMHD1 was added into the chamber to final concentration of 300 nM. After additional 40 minutes the concentration of the dNTP inflow was reduced 10 fold to 1 μM either for all dNTPs (balanced reduction), or for all but one dNTP (imbalanced reduction).

When the experiment is performed with no SAMHD1 added, there is very little change in the dNTP concentration within the chamber over time, because passive depletion of the 10 mL chamber at the 100 μL/min flow is slow. In contrast, when SAMHD1 is added to an equimolar dNTP mixture, dNTP concentrations rapidly drop and then stabilize at lower levels that are different for different dNTPs but all within the 0.1-1 μM range. It is evident that the established dNTP concentrations are in dynamic equilibrium because dNTP levels re-equilibrate at values below 0.1 μM for all dNTPs when the dNTP concentrations of the inflow are reduced to 1 μM.

Results are very different when the experiments are performed with individual dNTPs. In agreement with the results described above, dCTP depletion by SAMHD1 at 10 μM starting concentration (Fig. 5H) is indistinguishable from the passive depletion without SAMHD1 present (Fig. 5H). Depletion of dTTP is also dramatically impaired compared to the equimolar dNTP mixture. These data confirm that pyrimidine depletion at physiological dNTP concentrations is critically dependent on purine availability. In contrast, depletion of dATP or dGTP is very similar in the mixture and for individual dNTPs. It is important to note that the SAMHD1-controlled dATP equilibration rate in these reactions (∼1.6e-3 sec^-1^) is almost 7 times higher than the dissociation rate of the dATP loaded SAMHD1 tetramer (*k*_*-1*_∼2.5e-4 sec^-1^), which further confirms that the dNTPase activity of the enzyme in these experiments is indeed controlled by the dNTP-dependent conformational changes within the tetramer and not by tetramer assembly/disassembly.

Additional evidence of facilitated dNTP depletion is offered by experiments in which the dNTP delivery into the reaction chamber is imbalanced (Fig. 5I). For example, delivery of excess dATP into the reaction volume results in further reduction in the dTTP and dCTP concentrations established in the SAMHD1-controlled dynamic equilibrium. Interestingly, dTTP excess also promotes more efficient depletion of dCTP. In contrast, dNTP depletion appears less robust when dCTP delivery into the reaction is elevated. dGTP excess has little effect on other dNTPs, which is another indication that dGTP may already dominate allosteric activation of SAMHD1 in an equimolar mixture. These findings further illustrate the intricate interdependence between the depletion and biosynthesis of different dNTPs.

## DISCUSSION

In this study, we show that the tetramerization-dependent activation of SAMHD1 encompasses two distinct sequential steps: tetramer assembly and tetramer activation. Both steps involve binding of allosteric dNTP ligands (Fig. 6). The initial assembly takes place with incomplete dNTP occupancy of the allosteric sites, whereas full loading of all A2 sites with dNTP ligands activates dNTP hydrolysis. Crystal structures of SAMHD1 fully loaded with allosteric ligands reveal a compact tetramer with 222 point-group symmetry, in which the allosteric ligands are sequestered at the interfaces between the subunits^44-46,48,49^. Our data imply that the partially loaded, inactive tetramer adopts a more open conformation, in which the vacant allosteric sites are accessible to the solvent. Extensive structural data suggest that SAMHD1 tetramerization involves dimerization of dimers, so the most likely interpretation of our findings is that the initial assembly of SAMHD1 tetramers from two dimers occurs upon the loading of two allosteric sites with dNTP ligands, whereas binding of two more dNTPs is required for tetramer activation. The transition between the inactive and active tetrameric states is accompanied by a conformational change that sequesters allosteric ligands and prevents their dissociation from the enzymatically active, compact SAMHD1 tetramer. Conversely, the dissociation of the enzymatically active SAMHD1 tetramer must also proceed through the partially loaded, inactive tetrameric intermediate.

**Figure 6.**
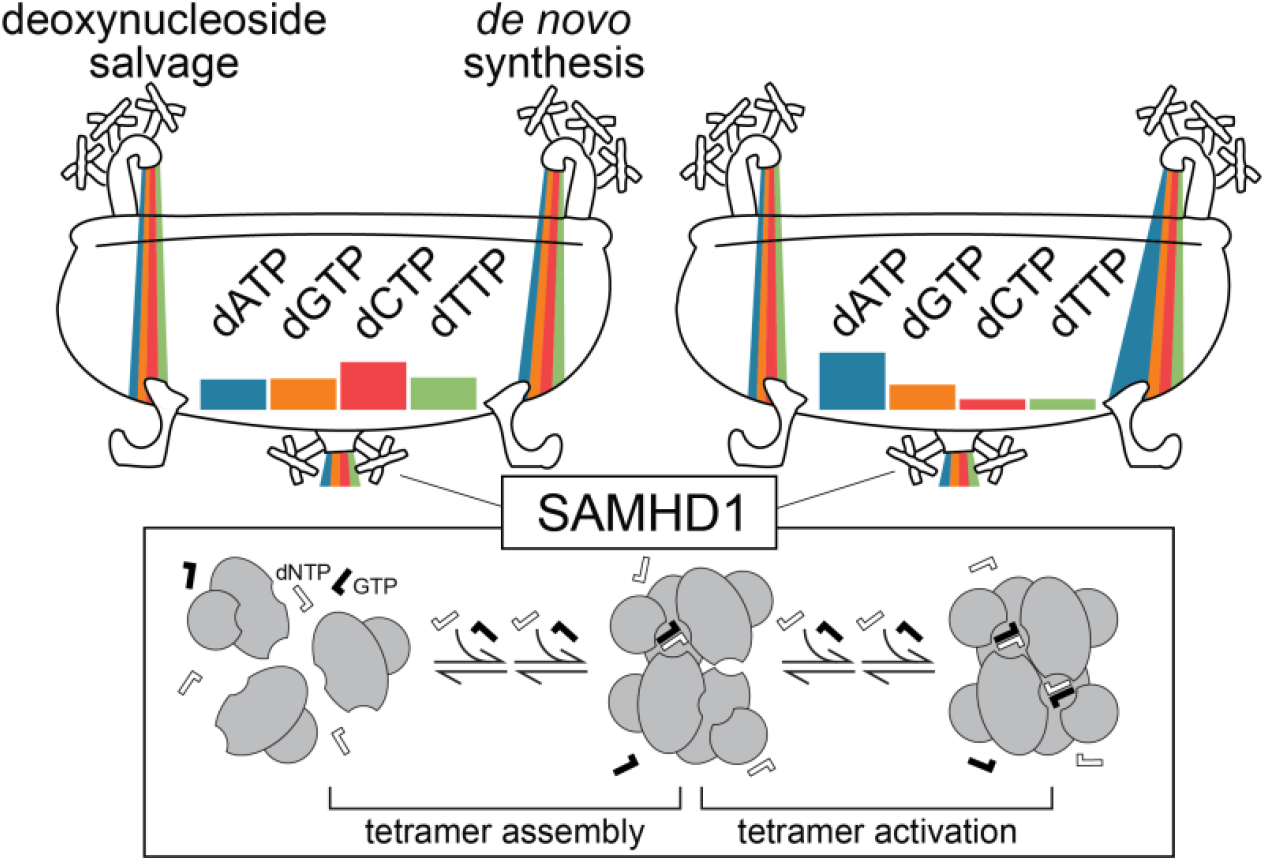
Allosteric activation of SAMHD1 shapes dNTP imbalances. Allosteric substrate activation mechanism consists of two sequential steps: tetramer assembly and tetramer activation (lower panel). The mechanism makes concentrations of different deoxynucleotides in the cellular dNTP pools interdependent and shapes the outcomes of SAMHD1-catalyzed dNTP depletion (upper panel). For example, the mechanism enables facilitated dNTP depletion by SAMHD1 and lowers equilibrium pyrimidine dNTP concentrations when production of dATP is elevated (upper right panel).

The equilibrium between the inactive and active tetrameric states is a key element in the SAMHD1-mediated regulation of dNTP homeostasis. The mechanism explains how SAMHD1 can circumvent the kinetic problem posed by the necessarily low association and dissociation rates of protein oligomerization. We show that the enzymatic activity is regulated by the binding and dissociation of allosteric dNTP ligands to a preformed SAMHD1 tetramer, which can happen much faster than the inherently limited assembly and disassembly of protein oligomers. Indeed, we demonstrate that the dynamic equilibration of dNTP concentrations by SAMHD1 is a faster process than tetramer disassembly.

The intermittent opening and closing of the tetramer, which is required for the binding and dissociation of allosteric dNTP ligands, provides for efficient sampling of the dNTP pools and shapes the outcomes of dNTP depletion by SAMHD1. The regulatory relationship between allosteric site occupancy and dNTP homeostasis has two integral parts. First, the overall catalytic efficiency of SAMHD1 tetramers depends on the identity of the four dNTP ligands occupying the allosteric sites. Second, the occupancy of the four sites is determined by the instantaneous dNTP concentrations and by the different affinities and cooperativities of allosteric dNTP binding. Differences in the affinity and cooperativity of dNTP binding to the allosteric sites that we report here explain and support conclusions reached in the pioneering structural analysis of SAMHD1 crystals formed in the presence of dNTP mixtures^44^. For example, strong negative cooperativity of dCTP binding explains why pure dCTP is a poor activator of SAMHD1, even though its ability to promote SAMHD1 tetramerization is comparable to that of other nucleotides. Notably, depletion of dNTPs is more robust in dNTP mixtures than for pure deoxynucleotides, which indicates that mixed loading of the allosteric sites with different dNTPs plays an important role in the regulation of dNTP levels. Better understanding of how the cooperativity of allosteric site loading with different dNTP combinations determines the outcomes of SAMHD1-mediated dNTP depletion will require further study.

A notable functional consequence of the allosteric activation mechanism is the phenomenon of facilitated dNTP depletion, which is manifested as the dependency of the equilibrium concentration of one nucleotide on the availability of another. This relationship is especially important for the regulation of pyrimidine dNTPs. Pyrimidines are less potent allosteric activators of SAMHD1, and their depletion critically depends on the availability of purine dNTPs. This functionality reveals how the outcomes of SAMHD1-mediated dNTP depletion are shaped by imbalances in dNTP biosynthesis by the de novo and salvage pathways (Fig. 6).

Our findings offer insight into the emerging functional roles of SAMHD1 in the biology of dNTP homeostasis. For example, facilitated dNTP depletion explains the recently reported antiviral effects of induced dNTP imbalances^58^. Supplementation of myeloid cells with a mixture of four deoxynucleosides abrogates the SAMHD1-mediated block to HIV-1 replication—an observation that strongly supports the role of the dNTPase activity in the restriction mechanism^5^. Remarkably, supplementation with a single nucleoside has the opposite effect on restriction. Elevated supply of a single deoxynucleotide lowers cellular concentrations of the other dNTPs and makes cells more restrictive to HIV-1 restriction^58^. The only exception is the supplementation with deoxycytosine, which has no effect on the availability of other deoxynucleotide triphosphates or on restriction. These observations are explained by the facilitated dNTP depletion functionality of SAMHD1 and the uniquely poor ability of dCTP to promote catalytic activation of SAMHD1. Notably, treatment of myeloid cells with Vpx results in a dramatic elevation of cellular purine dNTP pools, but only a modest increase in the dCTP and dTTP levels^59,60^. These findings suggest that dNTP biosynthesis rates are naturally imbalanced in myeloid cells and that facilitated dNTP depletion contributes to HIV-1 restriction even in the absence of the exogenously induced dNTP imbalances.

The mechanistic relationship between dNTP depletion and biosynthesis, as revealed by our study, explains why inhibitors of ribonucleotide reductase enhance the therapeutic efficacy of cytosine arabinoside (cytarabine or ara-C). Cytarabine is a nucleoside analog commonly used for the treatment of newly diagnosed and recurrent AML. SAMHD1 activity in AML cells is a critical determinant of cytarabine efficacy because the enzyme dephosphorylates the biologically active metabolite of cytarabine: cytarabine triphosphate^61,62^. Remarkably, inhibitors of ribonucleotide reductase suppress SAMHD1-mediated resistance to cytarabine^63^. The promise of RNR inhibition for improving outcomes of cytarabine-based therapies is currently being investigated in clinical trials^64^, but the molecular mechanism of this effect has remained obscure. Here we show that robust depletion of pyrimidine triphosphates by SAMHD1 depends on the availability of purine deoxynucleotide triphosphates, which explains why inhibition of ribonucleotide reductase impairs the ability of SAMHD1 to deplete cytosine arabinoside triphosphate.

The study also sheds light on the less well understood contributions of SAMHD1 to the regulation of dNTP homeostasis in proliferating cells. For example, the mechanism we describe offers unexpected insight into the emerging roles of SAMHD1 and dNTP metabolism in telomere biology^65,66^. One recent study focused on the genetics of telomere maintenance revealed that genetic disruption of thymidine biosynthesis enzymes caused significant shortening of telomeres, whereas SAMHD1 knockout had the opposite effect and markedly increased telomere length^66^. These observations suggest that facilitated depletion of thymidine by SAMHD1 controls thymidine availability at some specific point of the cell cycle and thus limits telomere length. The impact of SAMHD1 on telomere maintenance is intriguing, as it may contribute to the elevated occurrence of SAMHD1 mutations in cancer^67-70^.

Finally, a recent study of the pathology caused by mutations in purine nucleoside phosphorylase revealed that SAMHD1 activity limits T cell cytotoxicity resulting from the elevated rate of deoxyguanosine salvage^71^. The cytotoxicity of purine nucleoside phosphorylase mutations also depends on cytosine deaminase activity and dCTP availability. The findings support the existence of immune checkpoints controlled by relative concentrations of deoxynucleotides in the dNTP pools and the critical role of SAMHD1 in regulating these checkpoints.

In summary, the allosteric substrate activation of SAMHD1 reveals a mechanistic link between imbalances in dNTP biosynthesis and the outcomes of SAMHD1-mediated dNTP depletion (Fig. 6). The mechanism offers explanatory insight into the recently described contributions of SAMHD1 to the biology of dNTP homeostasis with implications for antiviral immunity, HIV/AIDS, telomere maintenance, therapeutic efficacy of nucleoside analogs and immune deficiencies caused by the imbalances in the dNTP supply.

## MATERIALS AND METHODS

### Protein Expression and Purification

The WT and mutant variants of the HD-domain SAMHD1_114-626_ constructs were cloned, expressed, purified using an *E. coli* expression system as described previously^17^. Briefly, SAMHD1_114-626_ constructs were cloned into pET30 vectors (Novagen) and transformed into *E. coli* BL21(DE3) cells. Bacterial cells were grown in LB media at 37 °C to an OD_600_ ∼ 0.6. SAMHD1 expression was induced by adding 1 mM isopropyl-B-D-thiogalactopyranoside followed by overnight incubation at 20 °C. Cells were harvested by centrifugation at 6000 x g for 15 minutes. Cells were then disrupted by sonication and cell lysate was centrifuged at 39,000 x g for 60 minutes. The supernatant was loaded onto a Strep-Tactin Sepharose (IBA Lifesciences) column equilibrated with 50 mM TRIS, pH 8, 1 M NaCl, and 20 mM B-mercaptoethanol. The protein was eluted with 2.5 mM desthiobiotin. The eluted protein was further purified on a Superdex 200 column (GE Healthcare) containing 50 mM TRIS, pH 8, 100 mM NaCl, and 1 mM TCEP. Purified proteins were stored at -80 °C in 20% glycerol.

### Sortase-catalyzed fluorescent labeling

N-terminal fluorescent labeling was carried out by sortase-catalyzed transpeptidation as described previously^72^. Briefly, SAMHD1 constructs were expressed with N-terminal affinity tags followed by a TEV protease cleavage site. Purified constructs were treated with TEV protease to remove the affinity tag leaving an N-terminal glycine residue on the cleaved protein. Short fluorescent peptides used for the N-terminal sortase-catalyzed labeling contained a fluorescent dye (chemically equivalent to either Alexa Fluor™> 488 (AF488) or Alexa Fluor™> 594 (AF594)) at the N-terminus and a sortase recognition sequence (KLPETGG) at the C-terminus. Labeling reactions were carried out using 50 μM fluorescently labeled peptide, 25 μM sortase, and 30 μM of the target protein for 45 minutes on ice. Unreacted fluorescent peptide was removed using the Superdex 75 size exclusion column (GE Healthcare) in 50 mM TRIS pH 8, 1 M NaCl and 1 mM TCEP. Labeling efficiency was calculated using sample absorbance at 280, 488 and 594 nm and the theoretical extinction coefficients of the protein and the dyes at these wavelengths. Samples used for FRET experiments had labeling efficiency >70%. Labeled proteins were stored at -80 °C in 20% glycerol.

### Förster resonance energy transfer (FRET) studies of SAMHD1 tetramerization

Förster resonance energy transfer (FRET) experiments were performed in 384-well plates (Corning 3575) using a Synergy 2 microplate reader (Biotek). Samples were prepared in FRET buffer containing 50 mM TRIS, pH 8, 100 mM NaCl, 5 mM MgSO_4_, 5 mM DTT, and 2 mg/mL BSA. All experiments were performed in a 40 μL final volume. Reactions, which contained equal concentrations of AF488-SAMHD1 and AF594-SAMHD1, were initiated by mixing fluorescently labeled proteins with varying concentrations of nucleotides and/or unlabeled protein as specified. Kinetic data were collected by repetitive measurements of three fluorescent intensity values (AF488 fluorescence intensity, AF594 fluorescence intensity, and FRET fluorescence intensity) over a specified time period for each sample. The following filter combinations were used for the three measurements: AF488 fluorescence intensity—485/20 nm excitation bandpass filter and 528/20 nm emission bandpass filter; AF594 fluorescence intensity—ex: 590/20 nm and em: 645/40 nm; FRET fluorescence intensity—ex: 485/20 nm and em: 645/40 nm. Normalized FRET intensity was calculated to minimize signal variations due to pipetting errors (Supplemental Methods). Tetramer dissociation rate datasets (Fig. 4E-H) were analyzed by numeric simulation and nonlinear least-squares fitting of the FRET decay data in MATLAB (Supplemental Methods).

### dNTPase assay with NMR detection

NMR dNTP assay was performed as described previously^54. 1^H NMR spectra were acquired on a Bruker 500-MHz spectrometer equipped with a 1.7-mm cryoprobe. Samples were prepared in a reaction buffer of 50 mM TRIS pH 7, 150 mM NaCl, 5 mM MgCl_2_, 5 mM DTT, 10% D_2_O. All reactions contained 1 μM SAMHD1, 500 μM of GTP, and the specified concentrations of one or more dNTP substrates in reaction buffer. Proton NMR spectra were recorded at regular 1-minute time intervals and the relative peak intensity of the proton signal of deoxynucleotide triphosphate (substrate) versus deoxynucleotide nucleoside (product) was measured as a function of time. *k*_*cat*_ and *K*_*m*_ values were determined by non-linear least-squares fitting of experimental progress curves to the Michaelis-Menten model in MATLAB as described in Supplemental Methods.

### dNTP quantification by LC-MS

Deoxynucleotide triphosphate concentrations were quantified by LC-MS on a Q Exactive Mass Spectrometer (ThermoFisher). dNTP chromatography was performed on a porous graphitic carbon column (Hypercarb; ThermoFisher) using a previously described strategy^73^. Glacial acetic acid was added to all dNTP samples to 10% final concentration and the samples were passed through a Nanosep 3k MWCO Omega spin membrane (PALL life sciences) to remove macromolecular contaminants. The membrane flowthrough was loaded onto a Hypercarb column (ThermoFisher) using a Dionex UltiMate 3000 HPLC and autosampler. The 20 μL samples were injected in mobile phase A (20 mM Ammonium Acetate, pH 9 in water) at a flow rate of 300 μL/min.

Nucleotides were eluted from the column with a 0% to 70% linear gradient of mobile phase B (100% acetonitrile) over a 5 minute time period. After the elution, the column was regenerated with 900 μL of mobile phase C (95% methanol) for each sample^73^. Mass spectra were recorded using four separate selective ion monitoring (SIM) scans in the negative ion mode targeting each of the four dNTPs: dATP SIM (488.4940 m/z -491.4940 m/z), dCTP SIM (464.4820 m/z – 467.4820 m/z), dGTP SIM (504.4880 m/z – 507.4880 m/z), dTTP SIM (479.4820 m/z – 482.4820 m/z). Peak volumes were integrated using Quant Browser in Xcalibur (Thermo Scientific).

### Continuous-flow studies of dNTP homeostasis

Studies of SAMHD1-controled dynamic equilibria of dNTP concentrations (Fig. 5) were performed using a 10 mL stirred-cell pressurized protein concentrator (Amicon; EMD Millipore) assembled with a 10 kDa MWCO nitrocellulose membrane (Millipore). The concentrator was pressurized with 50 psi of N_2_. The inflow rate was fixed at 100 μL/min and the outflow rate was adjusted to keep the cell volume constant at 10 ± 1 mL. The cell chamber buffer contained 50 mM TRIS pH 7, 150 mM NaCl, 5 mM MgSO_4_, 5 mM TCEP, and 2.5 mg/mL Bovine Hemoglobin (BHb). The inflow buffer contained 50 mM TRIS pH 7, 150 mM NaCl, 5 mM MgSO_4_, 5 mM TCEP, 50 μM GTP, and varying concentrations of dNTPs. SAMHD1 was added to the cell to 300 nM final concentration. dNTP concentrations in the outflow were measured by LC-MS as described above.

## ACKNOWLEDGEMENTS

This work was supported in part by NIH R01AI136697 (D.I.), Cancer Prevention and Research Institute of Texas (CPRIT) grant RP200058 (D.I.) and The Welch Foundation Research Grant AQ-1996-20190330 (D.I.). NMR and mass spectrometry analyses were conducted at the UTHSCSA Institutional Core Facilities supported in part by UTHSCSA and the Mays Cancer Center support grant NIH P30CA054174. We are grateful for the expert technical assistance of Sammy Pardo and Susan Weintraub.

## AUTHOR CONTRIBUTIONS

C.M. and C.H.Y. expressed and purified proteins, performed experiments, analyzed the data, prepared the figures, and co-wrote the manuscript. D.N.I. designed the study, prepared the figures, and wrote the manuscript with contributions from other co-authors.

## Declaration of generative AI and AI-assisted technologies in the editing process

During the preparation of this work the authors used GPT-4 to help identify typos and grammar/style issues in the final version of the manuscript. After using this tool/service, the authors reviewed and edited the content as needed and take full responsibility for the content of the publication.

