## Supplemental Information for "Allosteric substrate activation of SAMHD1 shapes deoxynucleotide triphosphate imbalances by interconnecting the depletion and biosynthesis of different dNTPs"

### SUPPLEMENTAL METHODS

#### Normalization of FRET data

To remove non-FRET sources of noise from the FRET datasets (e.g. pipetting errors, well bubbles, etc.) raw FRET intensity values were normalized using the following formula:

$$Normalized\ FRET = \frac{f_{FRET}[\,(\varepsilon 594)(f488) + (\varepsilon 488)(f594)]}{2(f488)(f594)}$$

Where,
$f_{FRET}$ = FRET fluorescence intensity
$f488$ = AF488 fluorescence intensity
$f594$ = AF594 fluorescence intensity
$\varepsilon 488$ = 488 nm extinction coefficient of the AF488-labeled protein (which takes into account labeling efficiency)
$\varepsilon 594$ = 594 nm extinction coefficient of the AF594-labeled protein (which takes into account labeling efficiency)

#### Analysis of FRET decay and dNTP depletion data in MATLAB

Modeling and analysis of the FRET decay (Fig. 4E-H) and dNTP depletion (Fig. 1 and Fig. 5) datasets was performed in MATLAB using the following general strategy: (1) theoretical progress curves were generated using

#### Modeling and analysis of the FRET decay and dNTP depletion datasets in MATLAB (Fig. 4E-H).

Theoretical progress curves (FRET vs time or dNTP vs time) were generated by numerically solving systems of differential equations (see below) derived from the model mechanism. The ODE45 nonstiff differential equation solver in MATLAB was used for this purpose. Values of model parameters were determined by nonlinear least-squares fitting of the theoretical progress curves generated by ODE45 to experimental datasets using the lsqcurvefit function in MATLAB.

In this study we demonstrate that SAMHD1 activation involves a tetrameric intermediate with incomplete loading of the A2 sites with allosteric dNTP ligands. The general model for such activation mechanism is shown in Fig. 4D:

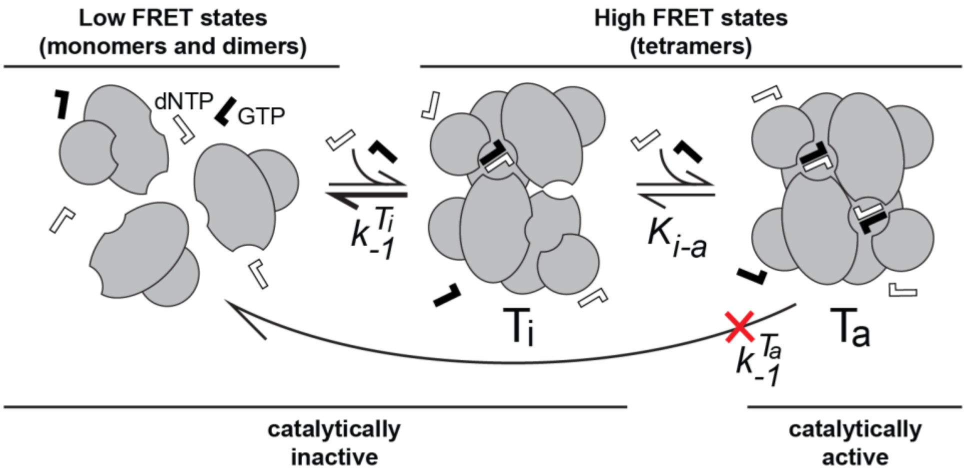


Based on this model, FRET decay datasets (Fig. 4B,C,F) can be described by the following system of two differential equations:

$$\begin{cases} \frac{\partial[dNTP]}{\partial t} = -0.5 * k_{cat}^{dNTP} * \left( \left(E_a + [dNTP] + K_m^{dNTP}\right) - \sqrt{\left(E_a + [dNTP] + K_m^{dNTP}\right)^2 - 4 * E_a * [dNTP]} \right) \\ \frac{\partial[T_t]}{\partial t} = -k_{-1}^{T_i} * T_i - k_{-1}^{T_a} * T_a \end{cases}$$

Where:
The first equation describes time dependence of the dNTP concentrations as dNTPs are being hydrolyzed by the WT SAMHD1. The second equation describes gradual dissociation of high-FRET SAMHD1 tetramers and the resulting decay of the FRET signal over time.

$T_t$ – is the fraction of total protein that exists in all tetrameric states; $0 \leq T_t \leq 1$
dNTP concentration at the beginning of each FRET decay experiment is significantly larger than the $EC_{50}^{dNTP}$ :

$$[dNTP]_{t=0} \gg EC_{50}^{dNTP},$$

therefore, one can assume that almost all protein is tetrameric:

$$T_t \approx 1$$

When the FRET signal intensity is normalized to decay from 1 to 0, then FRET=$T_t$ at any time point.

In the model in the Figure 4D the high-FRET states consist of the fully-dNTP-loaded, catalytically active tetramer denoted $T_a$ and the partially loaded tetrameric intermediate $T_i$:

$$T_t = T_i + T_a$$

The decay of the FRET signal over time is described by the second equation in the system above, where

$$\frac{\partial[T_t]}{\partial t} = -k_{-1}^{T_i} * T_i - k_{-1}^{T_a} * T_a$$

Data shown in Figure 4BC demonstrate that dissociation of $T_a$ is very slow. Therefore, tetramer dissociation and FRET decay can be approximated by:

$$\frac{\partial[T_t]}{\partial t} = -k_{-1}^{T_i} * T_i$$

When the equilibrium between the $T_i$ and $T_a$ is modelled using the Hill equation, and the equilibrium is assumed to be established faster than the $k_{-1}^{T_i}$ , then at any time point:

$$T_i = \frac{T_t * (K_{i-a})^n}{((K_{i-a})^n + [dNTP]^n)} \text{ and } T_a = \frac{T_t * [dNTP]^n}{((K_{i-a})^n + [dNTP]^n)}$$

where $K_{i-a}$ is the equilibrium constant and n is the Hill coefficient.

$E_a$ in the first equation is the concentration of the enzymatically active protein. If one assumes that only the $T_a$ state is catalytically active, then $E_a = [WT\ SAMHD1]_{total} * T_a$. In contrast, if one assumes that all tetrameric states are equally active, then $E_a = [WT\ SAMHD1]_{total} * T_t$. The model predicts very different dNTP depletion outcomes for these two scenarios (Supplemental Fig. S4B). Collectively, data presented in this study demonstrate that the dNTPase activity is upregulated by full loading of the A2 sites with dNTP ligands and that the former of the two assumptions is correct:

$$E_a = [WT\ SAMHD1]_{total} * T_a$$

Theoretical predictions of the FRET decay and dNTP depletion progress curves generated by the model above are shown in Supplementary Fig. S3D and Supplementary Fig. S4B, respectively.

If one assumes that the equilibrium between the inactive and active tetrameric states is accompanied by binding of two dNTP ligands (see Results Fig. 4G), the activation pathway can be described as follows:

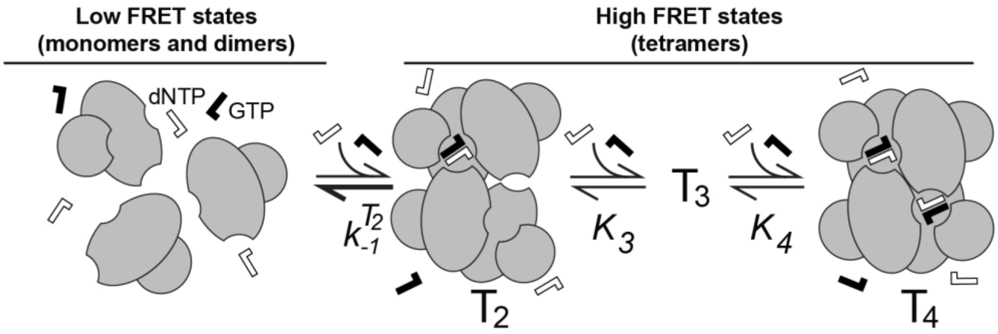


In this model,

$$T_t = T_2 + T_3 + T_4$$

where the subscripts denote the number of allosteric dNTP ligands bound to the three tetrameric states. Relative concentrations of the three tetrameric states at any time point can then be calculated using the Adair equation:

$$T_2 = \frac{T_t}{\left(1+\frac{[dNTP]}{K_3}+\frac{[dNTP]^2}{K_3K_4}\right)};\ T_3 = \frac{T_t*\frac{[dNTP]}{K_3}}{\left(1+\frac{[dNTP]}{K_3}+\frac{[dNTP]^2}{K_3K_4}\right)};\ T_4 = \frac{T_t*\frac{[dNTP]^2}{K_3K_4}}{\left(1+\frac{[dNTP]}{K_3}+\frac{[dNTP]^2}{K_3K_4}\right)}$$

In this model we also assume that only the fully dNTP loaded tetramer is catalytically active:

$$E_a = [WT\ SAMHD1]_{total} * T_4,$$

and that tetramer dissociation must proceed through the partially loaded tetrameric intermediate $T_2$

$$\frac{\partial[T_t]}{\partial t} = -k_{-1}^{T_2} * T_2$$

Values of model parameters $k_{-1}^{T_2}$, $K_3$ and $K_4$ listed in the Fig. 4G, were determined by least-squares fitting of the experimental data in Fig. 4F to the Adair model described above using the lsqcurvefit function in MATLAB.

**NMR-monitored dNTPase data analysis (Fig. 2B-E)**

$k_{cat}$ and $K_m$ values for each dNTP substrate were determined from the dNTPase progress curve data using a similar strategy to the one described in the previous section. Theoretical progress curves were calculated from the differential equations describing Michaelis-Menten kinetics (see below) using the ODE45 solver in MATLAB. $k_{cat}$ and $K_m$ values were determined by nonlinear least-squares fitting of the theoretical progress curves to experimental data using the lsqcurvefit function in MATLAB.

Analysis of single-substrate reactions was performed using the standard Michaelis-Menten equation:

$$\frac{\partial[dNTP]}{\partial t} = -1 * k_{cat}^{dNTP} * [SAMHD1] * \left(\frac{[dNTP]}{K_m^{dNTP} + [dNTP]}\right)$$

Reactions containing multiple substrates were analyzed by solving the following system of four equations:

$$\frac{\partial[dATP]}{\partial t} = -1 * k_{cat}^{dATP} * [SAMHD1] * \left(\frac{[dATP]}{K_m^{dATP}\left(1+\frac{[dATP]}{K_m^{dATP}}+\frac{[dGTP]}{K_m^{dGTP}}+\frac{[dCTP]}{K_m^{dCTP}}+\frac{[dTTP]}{K_m^{dTTP}}\right)}\right)$$

$$\frac{\partial[dGTP]}{\partial t} = -1 * k_{cat}^{dGTP} * [SAMHD1] * \left(\frac{[dGTP]}{K_m^{dGTP}\left(1+\frac{[dATP]}{K_m^{dATP}}+\frac{[dGTP]}{K_m^{dGTP}}+\frac{[dCTP]}{K_m^{dCTP}}+\frac{[dTTP]}{K_m^{dTTP}}\right)}\right)$$

$$\frac{\partial[dCTP]}{\partial t} = -1 * k_{cat}^{dCTP} * [SAMHD1] * \left( \frac{[dCTP]}{K_m^{dCTP} \left( 1 + \frac{[dATP]}{K_m^{dATP}} + \frac{[dGTP]}{K_m^{dGTP}} + \frac{[dCTP]}{K_m^{dCTP}} + \frac{[dTTP]}{K_m^{dTTP}} \right)} \right)$$

$$\frac{\partial[dTTP]}{\partial t} = -1 * k_{cat}^{dTTP} * [SAMHD1] * \left( \frac{[dTTP]}{K_m^{dTTP} \left( 1 + \frac{[dATP]}{K_m^{dATP}} + \frac{[dGTP]}{K_m^{dGTP}} + \frac{[dCTP]}{K_m^{dCTP}} + \frac{[dTTP]}{K_m^{dTTP}} \right)} \right)$$

For all dNTPs not included in the reaction, initial concentrations were set to zero: $[dNTP]_{t=0} = 0$

### SUPPLEMENTAL FIGURES

A

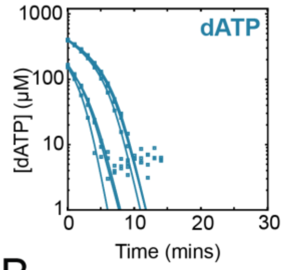


| dATP | | $k_{cat}$ ($s^{-1}$) | $K_m$ (μM) | $k_{cat}/K_m$ |
|---|---|---|---|---|
| T1 | 500 μM | 1.3 | 76 | 1.7E4 |
| | 250 μM | 1.3 | 89 | 1.5E4 |
| T2 | 500 μM | 1.1 | 70 | 1.7E4 |
| | 250 μM | 1.0 | 65 | 1.6E4 |
| T3 | 500 μM | 1.2 | 79 | 1.6E4 |
| | 250 μM | 1.3 | 64 | 2.0E4 |

B

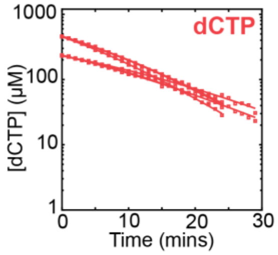


| dCTP | | $k_{cat}$ ($s^{-1}$) | $K_m$ (μM) | $k_{cat}/K_m$ |
|---|---|---|---|---|
| T1 | 500 μM | 1.6 | 690 | 2.3E3 |
| | 250 μM | 0.33 | 160 | 2.1E3 |
| T2 | 500 μM | 1.2 | 550 | 2.2E3 |
| | 250 μM | 0.56 | 350 | 1.6E3 |
| T3 | 500 μM | 1.3 | 620 | 2.1E3 |
| | 250 μM | 0.35 | 230 | 1.6E3 |

C

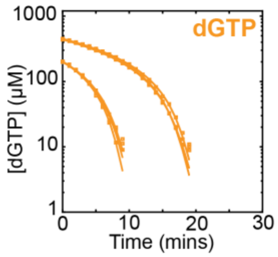


| dGTP | | $k_{cat}$ ($s^{-1}$) | $K_m$ (μM) | $k_{cat}/K_m$ |
|---|---|---|---|---|
| T1 | 500 μM | 0.51 | 33 | 1.5E4 |
| | 250 μM | 0.56 | 27 | 2.1E4 |
| T2 | 500 μM | 0.51 | 29 | 1.8E4 |
| | 250 μM | 0.52 | 28 | 1.8E4 |
| T3 | 500 μM | 0.49 | 26 | 1.9E4 |
| | 250 μM | 0.52 | 26 | 2.0E4 |

D

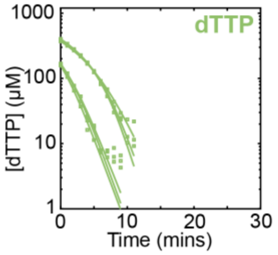


| dTTP | | $k_{cat}$ ($s^{-1}$) | $K_m$ (μM) | $k_{cat}/K_m$ |
|---|---|---|---|---|
| T1 | 500 μM | 1.3 | 100 | 1.2E4 |
| | 250 μM | 1.6 | 130 | 1.2E4 |
| T2 | 500 μM | 1.4 | 120 | 1.1E4 |
| | 250 μM | 2.0 | 200 | 1.0E4 |
| T3 | 500 μM | 1.4 | 170 | 8.4E3 |
| | 250 μM | 2.7 | 260 | 1.0E4 |

E

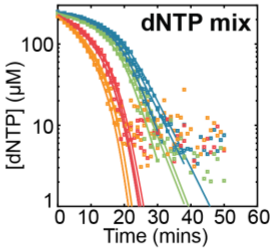


| dATP | | $k_{cat}$ ($s^{-1}$) | $K_m$ (μM) | $k_{cat}/K_m$ |
|---|---|---|---|---|
| T1 | 250 μM | 2.3 | 1000 | 2.3E3 |
| T2 | 250 μM | 1.8 | 440 | 4.0E3 |
| T3 | 250 μM | 2.7 | 610 | 4.4E3 |

| dCTP | | $k_{cat}$ ($s^{-1}$) | $K_m$ (μM) | $k_{cat}/K_m$ |
|---|---|---|---|---|
| T1 | 250 μM | 1.4 | 170 | 8.4E3 |
| T2 | 250 μM | 8.7 | 700 | 1.2E4 |
| T3 | 250 μM | 1.5 | 120 | 1.2E4 |

| dGTP | | $k_{cat}$ ($s^{-1}$) | $K_m$ (μM) | $k_{cat}/K_m$ |
|---|---|---|---|---|
| T1 | 250 μM | 0.21 | 16 | 1.3E4 |
| T2 | 250 μM | 0.35 | 19 | 1.8E4 |
| T3 | 250 μM | 0.48 | 28 | 1.7E4 |

| dTTP | | $k_{cat}$ ($s^{-1}$) | $K_m$ (μM) | $k_{cat}/K_m$ |
|---|---|---|---|---|
| T1 | 250 μM | 0.36 | 110 | 3.2E3 |
| T2 | 250 μM | 0.64 | 120 | 5.1E3 |
| T3 | 250 μM | 0.73 | 130 | 5.4E3 |

**Supplemental Figure S1: NMR dNTP depletion replicates of individual versus mixed dNTP depletion by NMR dNTPase assay.** (A-D) Replicates of individual dNTP depletion at two different substrate concentrations with calculated $k_{cat}$ and $K_m$ values for (A) dATP, (B) dCTP, (C) dGTP, and (D) dTTP substrates. (E) Replicates of equimolar dNTP mix with calculated $k_{cat}$ and $K_m$ values.

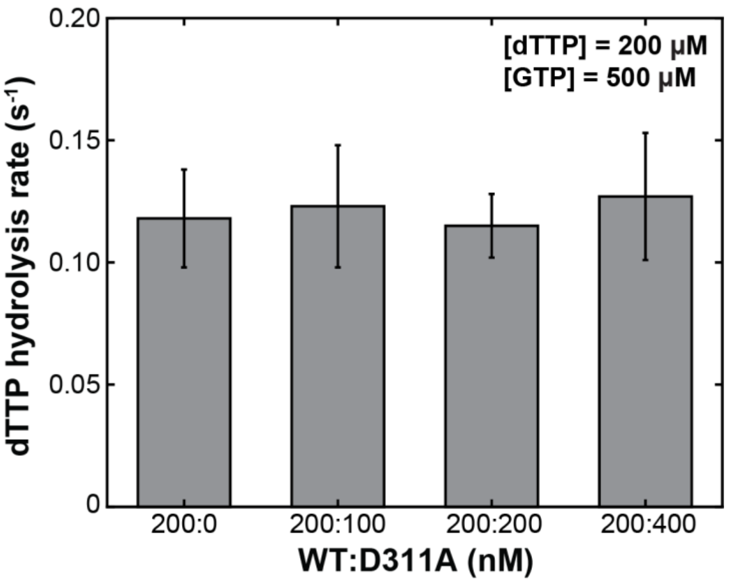


**Supplemental Figure S2: The catalytic activity of SAMHD1 tetramers is only dependent on the amount of wild-type enzyme present.** NMR dNTPase assay monitoring dTTP depletion by 200 nM WT SAMHD1 with varying concentrations of SAMHD1 D311A mutant added.

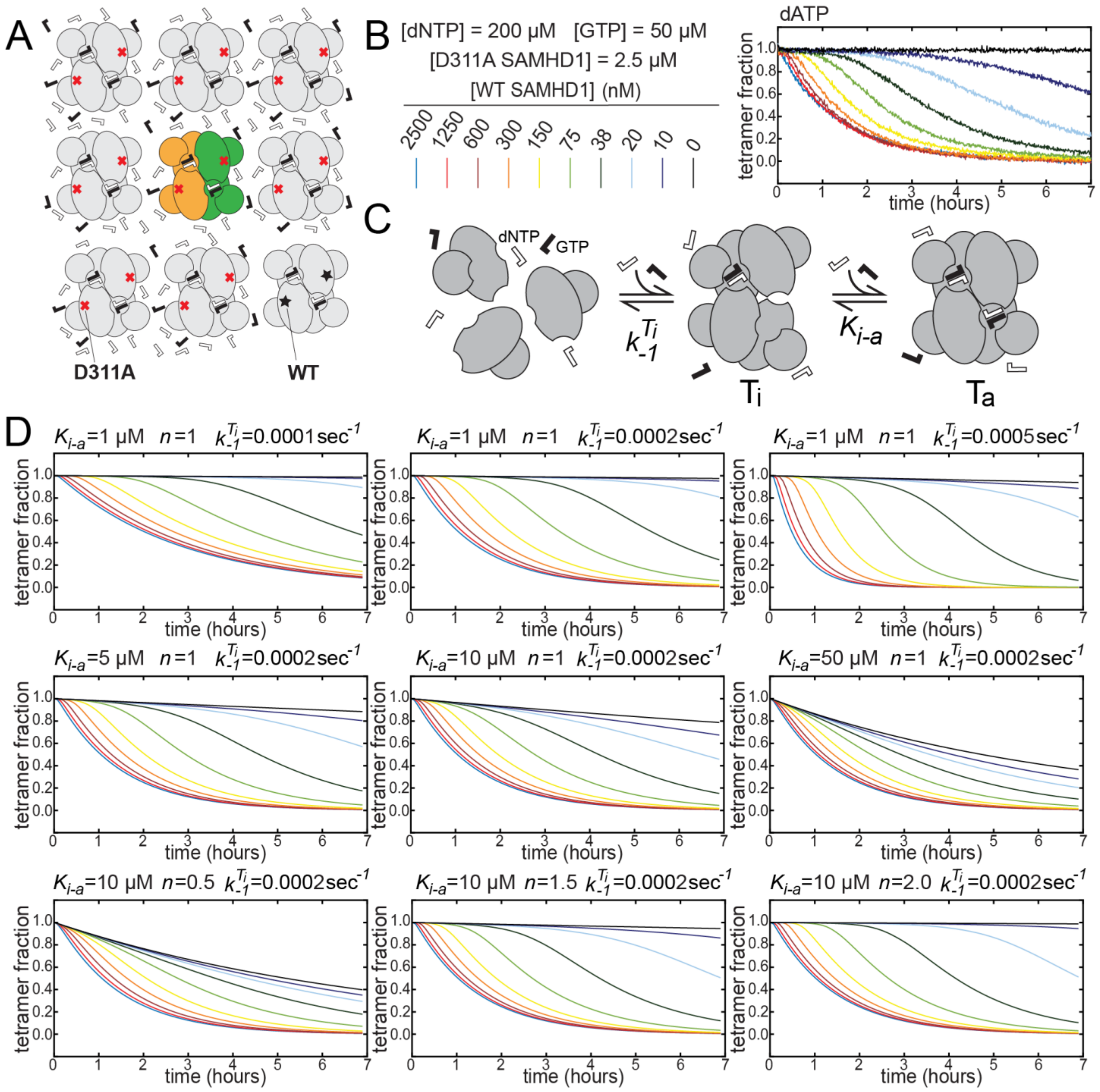


**Supplemental Figure S3. Modeling of FRET decay data.** (A) FRET decay experiments were performed by diluting preassembled high-FRET D311A tetramers into a mixture containing 2.5 μM unlabeled D311A SAMHD1 and increasing amounts of catalytically active WT SAMHD1. (B) Experimental dATP FRET decay dataset with individual traces for different WT SAMHD1 concentrations shown in different colors. (C) SAMHD1 activation model, in which the equilibrium between the $T_i$ and $T_a$ states is described by the Hill equation with the ligand concentration producing half occupation $K_{i\text{-}a}$ and the Hill coefficient $n$. (D) Theoretical FRET decay curves predicted by the model for different values of model parameters: $T_i$ dissociation rate $k_{-1}^{T_i}$ (top row), ligand concentration producing half occupation $K_{i\text{-}a}$ (middle row) and the Hill coefficient $n$ (bottom row).

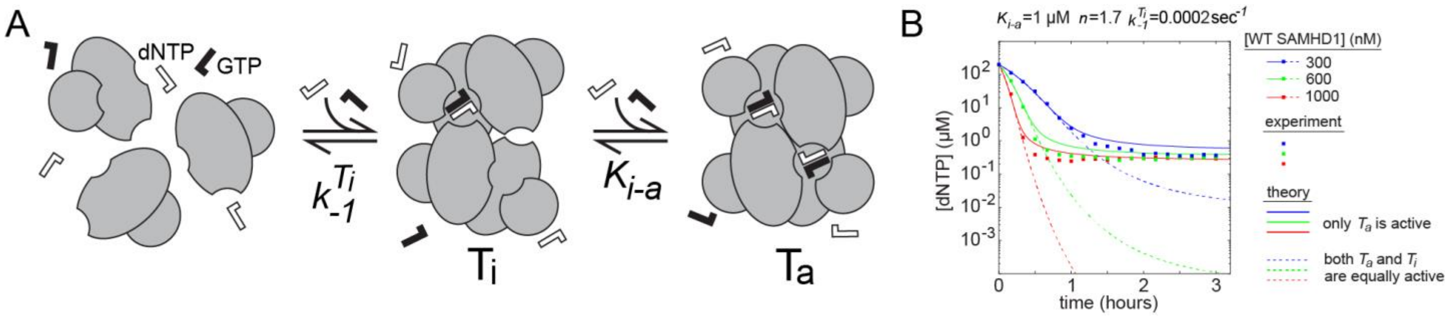


**Supplemental Figure S4. Modeling of dNTP depletion data.** (A) The SAMHD1 activation model can also be used to calculate theoretical dNTP depletion progress curves (see Supplemental Methods) (B) Owing to the relatively slow $T_i$ dissociation rate $k_{-1}^{T_i}$, the model predicts very different dNTP depletion outcomes depending on whether only the $T_a$ state is active (solid lines), or whether $T_a$ and $T_i$ are equally active (dashed lines). Experimental data (squares) demonstrates that $T_a$ is much more active than $T_i$ and, therefore, the dNTPase activity of SAMHD1 is controlled by the equilibrium between the $T_a$ and $T_i$ states.
